# Regularized Bagged Canonical Component Analysis for Multiclass Learning in Brain Imaging

**DOI:** 10.1101/698134

**Authors:** Carlos Sevilla-Salcedo, Vanessa Gómez-Verdejo, Jussi Tohka, Alzheimer’s Disease Neuroimaging Initiative

## Abstract

A fundamental problem of supervised learning algorithms for brain imaging applications is that the number of features far exceeds the number of subjects. In this paper, we propose a combined feature selection and extraction approach for multiclass problems. This method starts with a bagging procedure which calculates the sign consistency of the multivariate analysis (MVA) projection matrix feature-wise to determine the relevance of each feature. This relevance measure provides a parsimonious matrix, which is combined with a hypothesis test to automatically determine the number of selected features. Then, a novel MVA regularized with the sign and magnitude consistency of the features is used to generate a reduced set of summary components providing a compact data description.

We evaluated the proposed method with two multiclass brain imaging problems: 1) the classification of the elderly subjects in four classes (cognitively normal, stable mild cognitive impairment (MCI), MCI converting to AD in 3 years, and Alzheimer’s disease) based on structural brain imaging data from the ADNI cohort; 2) the classification of children in 3 classes (typically developing, and 2 types of Attention Deficit/Hyperactivity Disorder (ADHD)) based on functional connectivity. Experimental results confirmed that each brain image (defined by 29.852 features in the ADNI database and 61.425 in the ADHD) could be represented with only 30 – 45% of the original features. Furthermore, this information could be redefined into two or three summary components, providing not only a gain of interpretability but also classification rate improvements when compared to state-of-art reference methods.

## Introduction

Machine Learning (ML) techniques can be used for the design of imaging biomarkers for various brain disorders and, additionally, the inferred ML models can be analysed as multivariate, discriminative representations of the brain disease. Often in brain imaging applications of ML, the number of features is larger than the number of training subjects necessitating the use of dimensionality reduction techniques such as Feature Selection (FS) or Feature Extraction (FE). The usage of feature selection or extraction is critical in cases where the number of input variables is considerably greater than the number of data samples. In these cases, the usage of both methods entails dimensionality reduction and, subsequently, avoids overfitting problems.

For these reasons, there exist numerous studies proposing and applying dimensionality reduction methods in ML applications to brain imaging problems. The dimensionality reduction methods can be divided to three different categories, and combinations of them: 1) using a-priori neuroscientific information to select relevant features, for instance, volumes of certain regions of interest (Tanpitukpongse et al. 2017; Stoub et al. 2004; Douaud et al. 2013; Varon et al. 2015); 2) using unsupervised dimensionality reduction before the design of the classifier (e.g. Risacher et al. (2010); Klöppel et al. (2008); Hinrichs et al. (2011)) This dimensionality reduction is usually carried out with a principal component analysis (PCA). 3) Utilizing feature selection, either prior to the classifier design or jointly with the classifier design as a regularizer (e.g. Tohka et al. (2016); Michel et al. (2011); Cheng et al. (2017)).

Incidentally, while FS and supervised classification have been widely studied in brain imaging, the studies have focused on the binary classification, and multiclass setups have received only a limited amount of attention. The developed algorithms for multiclass classification in brain imaging are designed for specific problems (Bron et al. 2015; Qureshi et al. 2016; Yu et al. 2013) and can not be adapted to be used for other classification tasks than that they are designed for.^1^ Furthermore, most methods avoid dealing with high dimensional data and do not use FS or FE to automatically learn the relevant variables with the exception of regularized multinomial logistic regression that has found only few applications in brain imaging (Huttunen et al. 2013).

To address these shortcomings, we propose a Regularized Bagged - Cannonical Correlation Analysis (RB-CCA) method that is inspired by a recently proposed parsimonious Multivariate Analysis (pMVA) method for FE and FS (Muñoz-Romero et al. 2017). However, unlike with pMVA, the FS procedure of the RB-CCA is implemented by the calculation of a feature-wise sign consistency, analysed in Gomez-Verdejo et al. (2019), through a bagged Cannonical Correlation Analysis (CCA) approach. This, combined with a statistical test introduced in this paper assigning a p-value for the relevance of each feature, comprises an automatic feature selection method of the optimum characteristics for neuroimaging problems. The method’s goal is two-fold. First, it generates a parsimonious matrix (Nie et al. 2010; Chen and Huang 2012; Qureshi et al. 2016) which zeroes complete rows of the projection matrix and, thus, is capable of removing the irrelevant features. Second, the consistency of the feature is used to emphasize (by means of a proper regularization) the most relevant features of a subsequent CCA. This regularized CCA is capable of projecting the selected features onto a lower dimension space and, thus, providing a reduced subset of summary components to characterize the disease based on imaging data.

Furthermore, we propose the following novel contributions to pMVA to make it suitable for brain imaging tasks: (1) Class-wise feature selection to provide additional insights over the selected features; (2) A hypothesis test to automatically determine the number of selected features; (3) A dual formulation over the selected features to speed up the final feature extraction step; (4) A balanced version of the method to compensate the effect of class imbalance.

To analyse the performance of the proposed method, we use two different neuroimaging databases, Alzheimer’s Disease Neuroimaging Initiative (ADNI) and ADHD-200, Regarding the ADNI database we focus on classifying the subjects between 4 different groups: cognitively normal (NC), MCI subjects who will convert to AD within 3 years (progressive MCI), MCI subjects who do not convert to AD during 3 years (stable MCI), and subjects with AD. We will perform the classification using anatomical MRIs of the subjects, preprocessed using voxel-based morphometry (VBM). With the ADHD-200 database, we focus on the classifying the subjects between 3 groups: Typically Developing Children (TDC), ADHD of Inattentive type (ADHD-I) and ADHD of Combined type (ADHD-C). We perform the classification using functional connectivity measures between brain regions, extracted based on resting state functional MRI. The intuition with both databases is that the inclusion of all classes provide correlated and complementary information about the brain disorders in question (Li et al. 2016).

To demonstrate the advantages of the proposed method, its performance has been measured in comparison to various baseline methods: a linear SVM, to obtain a reference error without any dimensionality reduction, a standard CCA, to show the limitations of a feature extraction on its own, the feature selection (RFE) and classifying (HELM) methods analysed by Qureshi et al. (2016) and the feature selection method proposed by Abdulkadir et al. (2014), to analyse the advantages of our feature selector. We show that the proposed method outperforms the baseline methods in the classification accuracy.

## Methods: Regularized Bagged - CCA for Multiclass Learning

This section presents the Regularized Bagged - CCA (RB-CCA) method. As shown in the diagram in Figure 1, the method consists of two main steps:

1. **Feature selection** process. This first step combines a standard CCA with a bagging procedure to obtain a subset of selected voxels (*X*_*S*_) together with a measure of the relevance for each selected feature (*ρ*).
2. **Feature extraction for summary components design** to characterize each subject. This second step is based on a regularized version of CCA, guided by the variable relevance *ρ*, to reduce the input set of selected features to a subset of summary components (*X*_*B*_).

**Fig. 1:**
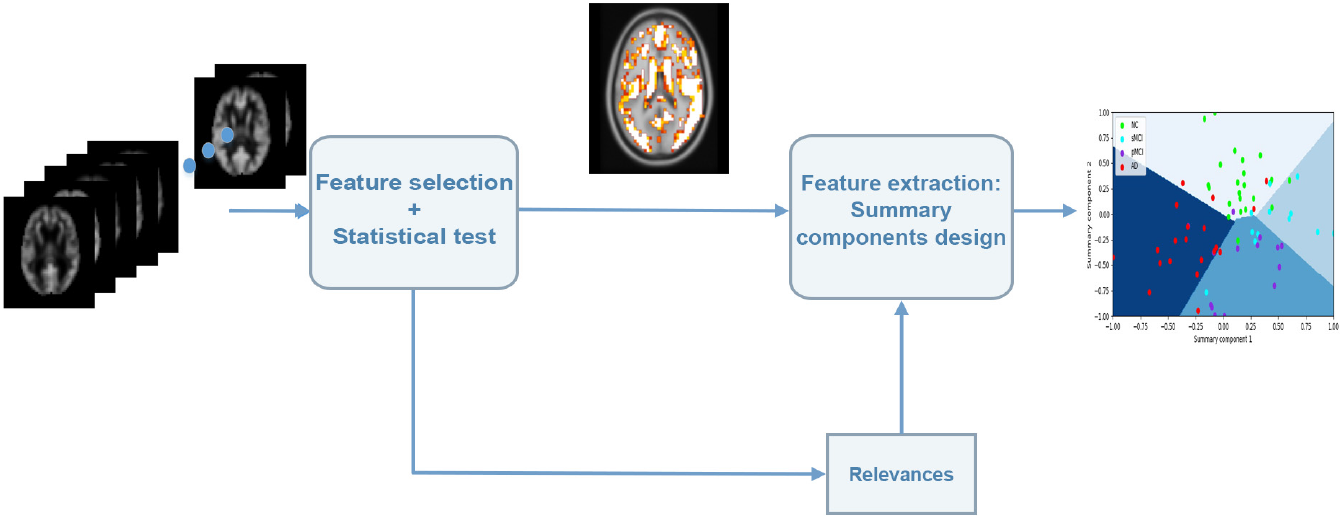
RB-CCA scheme for neuromarkers design.

### Review of the MVA framework

This section reviews the generalized MVA formulation presented in Muñoz-Romero et al. (2017, 2016), which unifies into a single framework the formulations of the most well-known MVA methods: PCA, CCA and OPLS.

In this context, a ML problem is given by N input/output data pairs 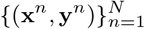, where the observations x^*n*^ have *d* features and the targets y^*n*^ have *c* output variables. Therefore, the problem is defined by two matrices: an input data matrix *X* ∈ ℝ^*N*×*d*^ and an output matrix *Y* ∈ ℝ^*N*×*c*^. In classification problems, this output matrix is encoded knowing 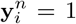 when x^*n*^ belongs to class *i* and 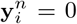 otherwise. In this work, we consider that the matrices *X* an *Y* are centred.

Our goal is to find a *d* × *R* projection matrix, *U*, that maps the input data onto a lower dimensional space with R features by solving the following optimization problem:

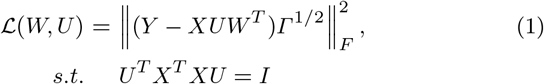

where 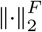 the Frobenius norm operator, *W* is a *c* × *R* regression matrix and *Γ* is the matrix that will define the different MVA algorithms considered: CCA (*Γ* = *N*(*Y*^*T*^*Y*)^−1^), PCA (*Γ* = *I* and *Y* = *X*) and OPLS (*Γ* = *I*).

The constraint in (1) can be replaced by one over *W*, obtaining an equivalent optimization problem:

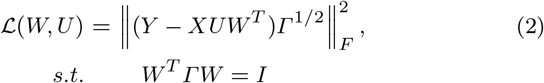

As this paper is focused on high dimensional small-sample problems (*d* >> *N*), working with the dual formulation results more computationally efficient algorithms. Noting that *U* can be expressed as a linear combination of the inputs and some dual variables *A*, *U* can be defined as *U* = *X*^*T*^*A* to express (2) as:

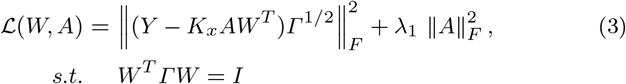

where we have defined *K*_*x*_ = *XX*^*T*^ as the linear kernel matrix of the input data and we have included a regularization term over *A* to overcome the ill-conditioned problems.

To solve our MVA problem, we firstly express *A* as a function of *W*^2^:

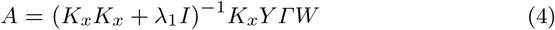

and, then, we substitute (4) into (3). Finally, *W* is obtained as the solution of the following eigenvalue problem:

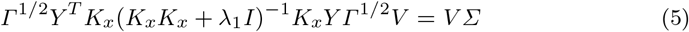

where *V* = *Γ*^1/2^*W* is introduced to simplify the computations. In a similar way, *V* can be computed and used to calculate *A*:

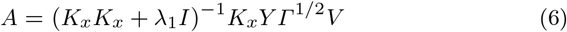

Note that the solution of Equation (5) involves operating with matrices of size of the order of *c*, instead of classical MVA approaches which work with matrices of size of the order of *N*. This advantage signifies a reduction of computational cost in almost all cases as the number of classes in a classification problem are usually considerably lower than the number of training data (*c* << *N*).

### Bagged MVA for feature selection

When a MVA method is applied, we obtain a new low-dimensional representation of the data given by

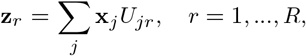

where *R* is the number of principal components found by the MVA method. Intuitively, one could analyse matrix *U* to measure the relevance of each characteristic according to the magnitude of the associated weights and, therefore, to generate a feature selector; however, in practice, this could cause overfitting problems when dealing with high dimensional problems. Therefore, as in Bi et al. (2003); Parrado-Hernández et al. (2014), overfitting can be mitigated by including of a bagging procedure (Breiman 2001).

Here, we propose to construct a set of *P* bagged MVA, where each MVA is trained with a randomly subsampled input data *X*_*M*_. This process is carried out class-wise, computing C sets of projection matrices for each bagging iteration, 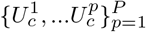.

Since the presented MVA framework works over the dual variables, the projection matrix in the dual space, *A*, can be calculated before the bagging procedure, as proposed in Muñoz-Romero et al. (2017). At the same time, using the labels corresponding to each subject, matrices *X* and *A* can be divided into the different classes, *X*_*c*_ and *A*_*c*_, where these matrices are defined by the rows of *X* and *A* corresponding to class *c*. This way the FS is able to work class-wise, having a more informative selection of the most relevant characteristics. Finally, these matrices can be randomly subsampled for the bagging procedure, and then, by calculating the product of both matrices 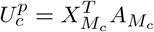, with *c* = 1,…,*C*, we can obtain the projection matrix for each class. This approach is fast, needing just to iterate a single matrix-product per class, which is a low cost operation.

Once the projection matrices, 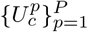 are computed, we can analyse their sign consistency to measure the relevance of each input feature for each eigenvector over the c-th class as:

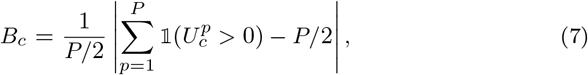

where 𝟙(T > 0) is the indicator function, which assigns a 1 to all the positive values in the matrix and, conversely, a 0 to the negative values. This new *d* × *R* matrix provides a high value when the j-th feature is sign consistent in the k-th eigenvector over the bagging iterations for the c-th class. This measures can be converted into a single measure for each feature and class by calculating their averaged value over the different eigenvectors:

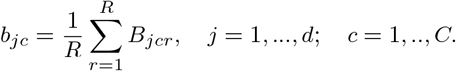

Note that the value of *b*_*jc*_ is normalized, so a value close to 1 implies a highly consistent feature, whilst a value close to 0 implies a non relevant feature with no consistency. By sorting the *b*_*jc*_ values, we have the class-wise most relevant features and can choose the number of those features we want to use. This selection could be carried out by adjusting the percentage of most relevant features (or selecting a threshold) by CV process. To avoid the computational cost of this process, next subsection introduces a hypothesis test to automatically fix the number of selected features. The scheme of this approach, in combination with the statistical test, is presented in Figure 2.

**Fig. 2:**
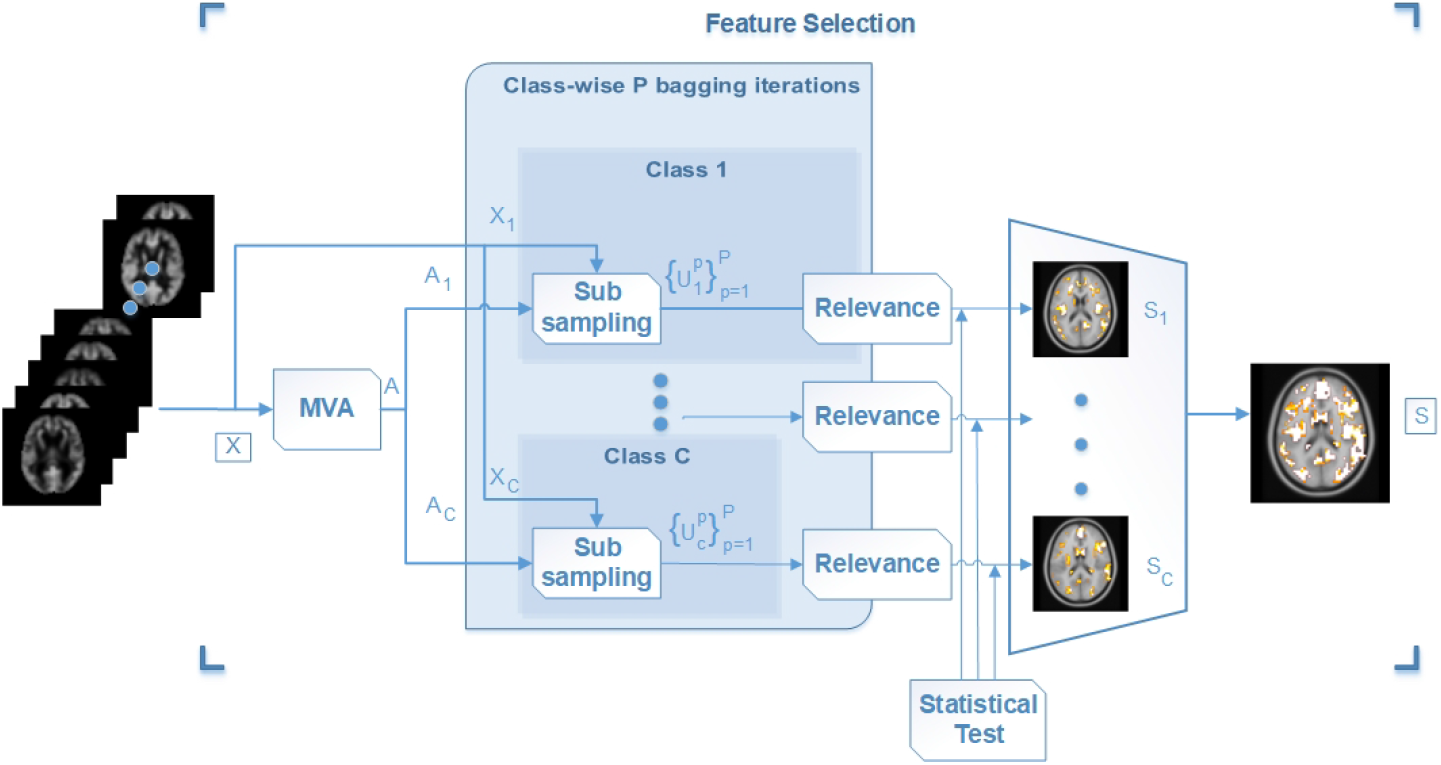
Class-wise feature Selection scheme for parsimonious MVA.

### Hypothesis test for feature selection

After applying the bagged MVA approach, a variable is irrelevant when it has positive and negative signs with the same probability. To be more precise, a variable *j* can be considered as irrelevant for the class c and the eigenvector r if its associated success probability

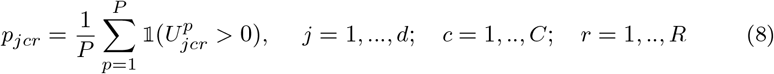

is equal to 0.5. Then, we can formulate the following hypothesis test:

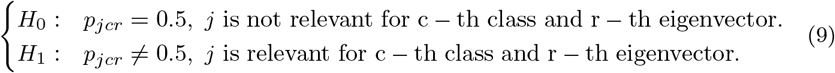

To statistically evaluate if *p*_*jcr*_ differs from 0.5, we define the statistic *t*_*jcr*_ which is given by the success probability divided by a scaling factor associated with the standard deviation of the probability. The derivation of this scaling factor is presented in A:

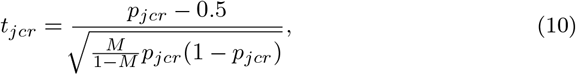

where *M* is the subsampling rate.

Under the null hypothesis, the statistic *t*_*jcr*_ follows a normal distribution with zero mean and unit standard deviation. Therefore, we can apply the test by selecting the values that correspond to the tails of the normal distribution.

Once the statistic *t*_*jcr*_ is computed, the class-wise feature selection can be determined by majority vote of the r parameter. This way, the selection takes into account if a feature is relevant for most eigenvectors or just for some of them.

Thanks to the inclusion of the statistical test, the cross-validation (CV) of the optimum number of selected features is not needed reducing the computational time. Furthermore, this efficient approach allows the selection of features in a class-wise manner, improving the interpretability of the results and posing an advantage over the approach presented in Muñoz-Romero et al. (2017).

### Regularized MVA

This section introduces the final step of the proposed method that, combining the parsimonious pattern and the relevance of each feature (learned in the previous step), will allow to compute the desired summary components.

Once the bagged MVA-based feature selection (BagMVA-ST) is applied, we obtain a parsimonious pattern defined by sets of indexes *S*_*c*_, with *c* = 1,…,*C*, which indicate which features are relevant for each class. These sets of indexes are then reduced to a single set of selected features, *S*, composed by the union of all *S*_*c*_ subsets. So, from the original data matrix *X*, here, we will use as input for this stage the matrix *X*_*S*_ consisting of the columns indexed by *S*.

Moreover, from the bagging process, we have also obtained information about the relevance of each feature. Here, we will also use this information to regularize this MVA so that we can guide the summary component design with the relevance of each feature. In this way, this regularization will aim to assign lower (/higher) penalties to more (/less) relevant features, increasing (/reducing) their influence over the projected data. To define which features are considered more or less relevant, two criteria are combined:

– The sign consistency of the eigenvectors. As the bagged feature selection does, we can use the consistency values *b*_*jc*_ to evaluate the usefulness of a feature.
– The associated eigenvector magnitude. It is expected that the eigenvector weights associated to relevant variables have a greater value than useless ones. So we can reinforce the consistency values with the following measure of the magnitude the associated eigenvector components:

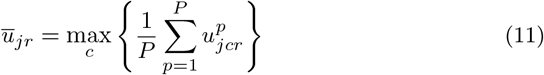

Then, we can combine both criteria to define the following relevance measure:

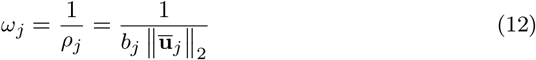

As high magnitude values imply more consistency, Equation (12) uses the inverse of the consistencies to indicate the relevancies for the regularization, in this way, more relevant features will have a low penalties and the regularized MVA will let them reach higher values.

Now, we can include this regularization over the MVA framework by replacing the penalty over dual variables of Equation (3) by a penalty over primal variables given by *ω*_*j*_ that is,

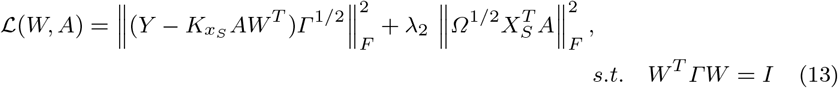

where 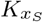 is the linear kernel matrix of the selected data, *Ω* is a diagonal matrix of the relevance measure values *ω*_*j*_ and *λ*_2_ is a regularization parameter. Despite feature selection, we are still dealing with a scenario where the input dimension (size of *S*) is still higher than *N*, so the dual space formulation leads to computational savings.

Then we can follow a similar procedure to that of Section Review of the MVA framework, so we can define the matrix *V* as *Γ*^1/2^*W* and obtain its value solving the following eigenvalue problem:

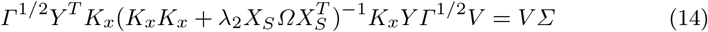

and, then, we can compute the solution of *A* by means of

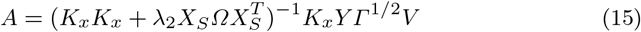

### Balanced regularized MVA

Neuroimaging problems can be highly imbalanced. The (class) imbalance refers here to the problem that there are different numbers of representative examples of each class, perhaps not reflecting the true (often unknown) class distribution. If we want to obtain a representative set of summary components (over all the classes), we have to make MVA approach pay more attention to the less populated classes.

For this purpose, we can define a new Frobenius norm as:

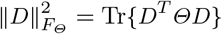

where *Θ* is a diagonal matrix which uses the values of Y to adjust weights inversely proportional to class frequencies in the input data as *N*/*N*_*c*_, being *N*_*c*_ the amount of samples of one class. Then, in order to balance the classes, (14) would be redefined as:

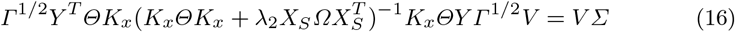

Conversely, with the inclusion of the class balance, (15) could be rewritten as:

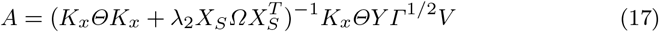

### Implementation details

We calculated the reported results with a nested 10-folds cross-validation. The outer CV is used to divide the dataset into training and test partitions, while the inner CV is in charge of validation and, therefore, it divides the training partition into a second training set and a validation set. This way we were able to estimate the performance of the whole framework and, additionally, validate the model parameters.

We used balanced classification accuracy of a one vs. all SVM to compare the performance of the different variations of the methods and to adjust the method hyperparameters. This balanced accuracy improves the performance of the methods on low-populated classes. The value of the hyperparameter *C* of the SVM was validated using a SVM with all the input voxels and was set to a rather small value (*C* = 0.035). We evaluated this value over the remaining approaches and we checked that their performance was good. Therefore, the parameter *C* of the SVM was set to *C* = 0.035 for all the methods under study, simplifying the CV of the MVA versions.

Despite the proposed framework includes several MVA approaches, for the sake of simplicity, we have limited the experimental comparison to the CCA approach. We made this decision based on the fact that CCA works in a supervised way and that it has been seen that CCA and OPLS work in a similar way in classification problems Sun et al. (2009).

We analysed the dependency on the subsampling rate and saw that the performance of the method does not depend on this value. Therefore, the subsampling rate was set to 50%. However, in order to have a balanced feature extractor and, learning the consistency of each input voxel equally over all the classes, the data was balanced in the bagging by randomly selecting the same number of samples in each class.

We cross-validated the regularization hyperparameter *λ*_2_, as its value was indeed critical for the final method’s performance. Its optimum value was cross-validated, exploring 17 values in a logarithmic scale from [10^−4^ to 10^3^]. At the same time, we set the number of extracted features to the maximum, *#classes* – 1, although some tests have been carried out to discard the relevance of using less features.

The implementation of this project was done using *Python 2.7.13* and the cross validation was carried out using the package *StratifiedKFold* from *Scikit-learn* (Pedregosa et al. 2011). An exemplary notebook, including the complete code of the proposed method, is available at https://github.com/sevisal/regMVA.git.

## Materials

### ADNI data

Data used in the preparation of this article were obtained from the Alzheimer’s Disease Neuroimaging Initiative (ADNI) database (adni.loni.usc.edu). The ADNI was launched in 2003 as a public-private partnership, led by Principal Investigator Michael W. Weiner, MD. The primary goal of ADNI has been to test whether serial magnetic resonance imaging (MRI), positron emission tomography (PET), other biological markers, and clinical and neuropsychological assessment can be combined to measure the progression of mild cognitive impairment (MCI) and early Alzheimer’s disease (AD).

The initial goal of ADNI (ADNI-1) was to recruit 800 subjects but ADNI has been followed by ADNI-GO and ADNI-2. To date these three protocols have recruited over 1500 adults, ages 55 to 90, to participate in the research, consisting of cognitively normal older individuals, people with early or late MCI, and people with early AD. The follow-up duration of each group is specified in the protocols for ADNI-1, ADNI-2 and ADNI-GO. For up-to-date information, see www.adni-info.org.

Data used in this work included MRIs from 200 AD patients, 164 pMCI subjects, 100 sMCI subjects and 231 NCs (T1-weighted MP-RAGE sequence at 1.5 T, typically 256 × 256 × 170 voxels with the voxel size of approximately 1*mm* × 1*mm* × 1.2*mm*) for whom baseline MRI data were available. The characteristics of these subjects are summarized in Table 1. The conversion status of the MCI subjects was defined as in Moradi et al. (2015). Briefly, a subject was considered to be in progressive MCI group if the diagnosis was MCI at the baseline and the subject converted to AD in three years. A subject was considered to be in stable MCI group if the diagnosis was MCI at the baseline and the subject did not convert to AD during the follow-up. Subjects who had less than 3 years of follow-up and subjects whose diagnostic status fluctuated were excluded.

**Table 1:** ADNI - Characteristic of data samples used in this work

| Characteristic | AD | pMCI | sMCI | NC |
| --- | --- | --- | --- | --- |
| No. of subjects | 200 | 164 | 100 | 231 |
| Age | $75.6 \pm 7.7$ | $74.57 \pm 7.0$ | $75.4 \pm 7.2$ | $76.0 \pm 5.0$ |
| Gender (M/F) | 103/97 | 97/67 | 66/34 | 119/112 |

The MRIs were preprocessed into gray matter tissue images in the stereostatic space, as described by Gaser et al. (2013), and thereafter they were smoothed with the 8-mm FWHM Gaussian kernel, resampled to 4 mm spatial resolution and masked into 29.852 voxels.

Atrophic regions detected in AD patients were found to overlap with those regions showing a normal age-related decline in healthy control subjects (Dukart et al. 2011). Therefore, the data was age-corrected by regressing out the age of the subject on a voxel-by-voxel basis (Moradi et al. 2015).

### ADHD data

We have also studied functional connectivity in ADHD using ADHD200 data (Milham et al. 2012). The data consists of 973 resting state fMRI and anatomical MRI datasets collected at eight independent imaging sites, all from children and adolescents between the ages of 7 and 21 years. We used the resting state fMRIs preprocessed by the Neuro Bureau using the Athena computer cluster of Virginia Tech as described by Bellec et al. (2017). Briefly, the preprocessing was done with AFNI (https://afni.nimh.nih.gov/) as detailed in https://www.nitrc.org/plugins/mwiki/index.php/neurobureau:AthenaPipeline#Extracted_Time_Courses.

The time courses of brain regions corresponding to CC400 atlas (ADHD-200 version, Craddock et al. (2012)) were obtained by averaging voxel-wise fMRI intensities within regions. This yielded 351 regional time courses per subject. Based on these 351 regional time courses, we computed a 351 × 351 correlation matrix describing the strength of the functional connectivity between region pairs. Vectorizing correlation matrix and removing redundant elements yields 61425 features per subject. We removed the datasets that did not pass the quality control of the Neuro Bureau. After this, data from 922 subjects remained (555 Typically Developing, 204 ADHD-Combined, 12 ADHD-Hyperactive/Impulsive, and 127 ADHD-Inattentive). We still removed 12 ADHD-Hyperactive/impulsive cases from consideration as the number of subjects in this group was not sufficient for meaningful classification.

## Results

### Performance compared to baseline methods

This section presents the experimental results obtained using the presented methods with both the ADNI and the ADHD databases. To analyse the performance of our algorithm, we have compared it with the following baseline approaches:

– SVM classifier (SVM): A set of original features are fed to a linear SVM.
– Standard CCA with a SVM (CCA): The original features are processed by a standard CCA and, later, classified by a linear SVM.
– SVM significance map with a SVM (p-map): The features are fed to the p-map of the SVM which carries out the feature selection, being after classified by a linear SVM. The p-map+SVM method was used by Abdulkadir et al. (2014) in the CADDementia challenge (Bron et al. 2015) as a multiclass classification approach following a one vs. all approach, providing a set of selected features for each class.

Regarding the ADHD database, we have implemented the methods used by Qureshi et al. (2016) for multiclass classification with feature selection. One should note that Qureshi et al. (2016) used 320 cortical features based on structural MRI whereas we have 61425 features based on resting-state fMRI. Three different implementations of the method have been used, using ELM with all the features (ELM), ELM along with the feature selection obtained with the RFE (ELM+RFE) and HELM with the feature selection of the RFE (HELM+RFE). When cross-validating the number of selected features, we used CV Stability Point (CV-SP) to select the optimum threshold^3^.

We compared the baseline approaches to the RB-CCA+ST (RB-CCA along with statistical test) version of our method with balanced CCA. We made use of our balancing MVA scheme (Subsection Balanced regularized MVA) along with the balanced version of the final SVM classifier. Different versions of our method are compared in section Analysis of the different stages of the method.

The results obtained with the different methods with the ADNI database are listed in Table 3. These show that the proposed approach outperformed the baseline methods both in terms of balanced accuracy and AUC. In addition, this performance improvement was achieved using one third of the original features. Comparing our approach with the p-map baseline, the proposed approach resulted in better classification accuracy with smaller standard deviation in the number of selected features and could thus be considered to lead to more consistent and relevant characterization of the classification problem.

**Table 2:** ADHD - Characteristic of data samples used in this work

| Characteristic | ADHD-C | ADHD-I | TDC |
| --- | --- | --- | --- |
| No. of subjects | 204 | 127 | 555 |
| Age | $11.94 \pm 2.6$ | $11.3 \pm 3.2$ | $12.2 \pm 3.5$ |
| Gender (M/F) | 170/34 | 92/35 | 288/267 |

**Table 3:** ADNI - Accuracy results with the proposed method compared with the baselines. This values have been obtained validating using the balanced accuracy. In this case, the balanced accuracy value obtained by chance would be 25%. The results show that the proposed method outperforms the baselines in terms of both accuracy and AUC.

| Method | No. feat | Accuracy | class AUC |  |  |  | AUC |
| --- | --- | --- | --- | --- | --- | --- | --- |
|  |  |  | NC | sMCI | pMCI | AD |  |
| SVM | 29.852 | 56,47<br>$\pm 5,97\%$ | 0,892 | 0,694 | 0,792 | 0,860 | 0,831<br>$\pm 0,024$ |
| Standard CCA | 29.852 | 52,15<br>$\pm 5,17\%$ | 0,854 | 0,603 | 0,751 | 0,825 | 0,786<br>$\pm 0,029$ |
| p-map | 19.637<br>$\pm 6.959$ | 55,98<br>$\pm 5,95\%$ | 0.886 | 0.694 | 0.795 | 0.862 | 0,830<br>$\pm 0,024$ |
| RB-CCA+ST<br>(Balanced) | 13.222<br>$\pm 1.004$ | 62,91<br>$\pm 4,64\%$ | 0,915 | 0,766 | 0,830 | 0,882 | 0,864<br>$\pm 0,024$ |

Table 4 shows the results obtained on the ADHD database. The studied methods improved the results that would be obtained by chance (33, 3% accuracy when randomly assigning a subject to one of the three possible classes). When comparing our method with the one proposed by Qureshi et al. (2016), our method outperformed it in both AUC and balanced accuracy term. In this dataset, the main advantages are in terms of interpretability, since we are capable of reducing the amount of input variables one fifth from the original, maintaining a good a classification score; note that the p-map approach presents poor performance results when reducing the number of input features.

**Table 4:** ADHD - Accuracy results with the proposed method compared with the baselines. This values have been obtained validating using the balanced accuracy. In this case, the balanced accuracy value obtained by chance would be 33, 3%. The results show that the proposed method performs in a similar way to the baselines, while been capable of reducing one fifth the amount of used voxels.

| Method | No. feat | Accuracy | class AUC |  |  | AUC |
| --- | --- | --- | --- | --- | --- | --- |
|  |  |  | TDC | ADHD-I | ADHD-C |  |
| SVM | 61.425 | 39,40<br>$\pm 14,66\%$ | 0,592 | 0,502 | 0,649 | 0,582<br>$\pm 0,057$ |
| Standard CCA | 61.425 | 36,94<br>$\pm 8,40\%$ | 0,597 | 0,500 | 0,632 | 0,581<br>$\pm 0,053$ |
| p-map | 969<br>$\pm 61$ | 36,54<br>$\pm 6,33\%$ | 0,568 | 0,484 | 0,635 | 0,568<br>$\pm 0,032$ |
| ELM | 61.425 | 24,26<br>$\pm 10,05\%$ | 0,533 | 0,504 | 0,567 | 0,537<br>$\pm 0,065$ |
| ELM + RFE | 7.025<br>$\pm 2.376$ | 22,64<br>$\pm 5,26\%$ | 0,507 | 0,511 | 0,534 | 0,514<br>$\pm 0,060$ |
| HELM + RFE | 7.025<br>$\pm 2.376$ | 28,99<br>$\pm 8,56\%$ | 0,525 | 0,505 | 0,564 | 0,531<br>$\pm 0,057$ |
| RB-CCA+ST<br>(Balanced) | 18.295<br>$\pm 4.393$ | 38,48<br>$\pm 8,18\%$ | 0,600 | 0,530 | 0,644 | 0,600<br>$\pm 0,058$ |

### Analysis of the different stages of the method

The proposed method combines different steps: a feature selection step, a statistical test and a regularized CCA. In this subsection, we will analyse the effect of these steps to the final performance of our method. To do so, we have included different combinations of the proposed feature selection (no FS, Bagged CCA with the statistical test based threshold (BagCCA+ST), Bagged CCA with the CV-based threshold (BagCCA+CV)) and extraction methods (no FE, standard CCA, regularized CCA). When using the CV-based thresholding, we used CV Stability Point to select the optimum threshold.

Table 5 depicts the accuracy results with the different versions of the method. All the methods in Table 5 are with balancing and the results of unbalanced method are presented in B. As is visible in the table, feature selection improved the performance. Between the two types of feature selection the statistical test based threshold slightly outperforms with respect to accuracy, while leading to substantial savings in computation time. Also, adding regularization to the CCA was beneficial.

**Table 5:** ADNI - Accuracies with the different versions of the method. This table justifies the need of adding the regularisation to the CCA as well as the usage of a selection method. The feature selection not only provides a better performance, but also improves the interpretability of the results. Furthermore, feature selecting with the statistical test is more efficient in terms of computational time than the validation.

| Extraction | Selection method | No. feat | Accuracy | class AUC |  |  |  | AUC |
| --- | --- | --- | --- | --- | --- | --- | --- | --- |
|  |  |  |  | NC | sMCI | pMCI | AD |  |
| No | None | 29.852 | 56,47<br>$\pm 5,97\%$ | 0,892 | 0,694 | 0,792 | 0,860 | 0,831<br>$\pm 0,024$ |
| | BagCCA+CV | 11.642<br>$\pm 5.373$ | 60,55<br>$\pm 6,48\%$ | 0,908 | 0,737 | 0,825 | 0,883 | 0,857<br>$\pm 0,026$ |
| | BagCCA+ST | 13.222<br>$\pm 1.004$ | 61,90<br>$\pm 9,65\%$ | 0,912 | 0,739 | 0,835 | 0,884 | 0,861<br>$\pm 0,024$ |
| Standard CCA | None | 29.852 | 52,09<br>$\pm 5,16\%$ | 0,854 | 0,593 | 0,747 | 0,825 | 0,783<br>$\pm 0,030$ |
| | BagCCA+CV | 15.075<br>$\pm 5.602$ | 52,12<br>$\pm 5,89\%$ | 0,850 | 0,598 | 0,745 | 0,831 | 0,783<br>$\pm 0,030$ |
| | BagCCA+ST | 13.222<br>$\pm 1.004$ | 52,07<br>$\pm 5,95\%$ | 0,856 | 0,635 | 0,754 | 0,828 | 0,792<br>$\pm 0,028$ |
| Regularized CCA | None | 29.852 | 58,19<br>$\pm 6,42\%$ | 0,911 | 0,718 | 0,816 | 0,870 | 0,849<br>$\pm 0,019$ |
| | BagCCA+CV | 6.862<br>$\pm 5.135$ | 62,04<br>$\pm 5,50\%$ | 0,909 | 0,785 | 0,835 | 0,887 | 0,868<br>$\pm 0,025$ |
| | BagCCA+ST | 13.222<br>$\pm 1.004$ | 62,91<br>$\pm 4,64\%$ | 0,915 | 0,766 | 0,830 | 0,882 | 0,864<br>$\pm 0,024$ |

Table 6 lists the results on the ADHD database with the different versions of the method. Comparing the selection methods BagCCA+ST and BagCCA+CV, the accuracy differences were minimal, however, BagCCA+ST was orders of magnitude faster due to the elimination of one CV loop. Furthermore, the number of selected features by the CV-SP had a greater standard deviation than by the ST meaning that the threshold selection by ST resulted in more stable feature selection. Feature selection using CV-SP, in some cases, was too conservative, selecting too few features.

**Table 6:** ADHD - Accuracy results with the different versions of the method, considering the usage of the proposed selection and extraction methods in their balanced version. The feature selection improves the interpretability of the results, reducing them by one fifth while keeping a similar performance. Furthermore, feature selecting with the statistical test is more efficient in terms of computational time than the validation.

| Extraction | Selection method | No. feat | Accuracy | class AUC |  |  | AUC |
| --- | --- | --- | --- | --- | --- | --- | --- |
|  |  |  |  | TDC | ADHD-I | ADHD-C |  |
| No | None | 61.425 | 39, 40<br>$\pm 14, 66\%$ | 0,592 | 0,502 | 0,649 | 0, 582<br>$\pm 0, 057$ |
| | BagCCA+CV | 21.806<br>$\pm 19.202$ | 36, 11<br>$\pm 10, 52\%$ | 0,564 | 0,501 | 0,634 | 0, 571<br>$\pm 0, 045$ |
| | BagCCA+ST | 18.295<br>$\pm 4.393$ | 38, 65<br>$\pm 12, 23\%$ | 0,589 | 0,516 | 0,634 | 0, 588<br>$\pm 0, 044$ |
| Standard CCA | None | 61.425 | 34, 53<br>$\pm 8, 78\%$ | 0,593 | 0,506 | 0,629 | 0, 589<br>$\pm 0, 057$ |
| | BagCCA+CV | 16.277<br>$\pm 11.978$ | 37, 18<br>$\pm 8, 72\%$ | 0,582 | 0,513 | 0,619 | 0, 580<br>$\pm 0, 051$ |
| | BagCCA+ST | 18.295<br>$\pm 4.393$ | 36, 18<br>$\pm 8, 96\%$ | 0,598 | 0,528 | 0,624 | 0, 594<br>$\pm 0, 039$ |
| Regularized CCA | None | 61.425 | 37, 10<br>$\pm 8, 73\%$ | 0,597 | 0,492 | 0,627 | 0, 589<br>$\pm 0, 055$ |
| | BagCCA+CV | 24.877<br>$\pm 17.879$ | 37, 40<br>$\pm 6, 96\%$ | 0,590 | 0,518 | 0,625 | 0, 587<br>$\pm 0, 050$ |
| | BagCCA+ST | 18.295<br>$\pm 4.393$ | 38, 48<br>$\pm 8, 18\%$ | 0,600 | 0,530 | 0,644 | 0, 600<br>$\pm 0, 058$ |

In conclusion, the usage of the regularized CCA with the bagged CCA and the statistical test (RB-CCA+ST) led to the best performance among the presented methods.

### Balanced accuracy in validation

Tables 7 and 8 show the confusion matrices of the classifiers obtained based on the balanced accuracy and standard accuracy, respectively. These were calculated as the sum of the confusion matrices over the 10 outer CV-folds. The validation is used to select the regularization parameter *λ*_2_ as outlined in section 2.6. If the balanced accuracy was used as the validation measure, the influence of the most populated classes was mitigated to give more importance to less populated classes. Instead, if the standard accuracy was used as validation measure, the influence of the more populated classes was not mitigated and, therefore, the misclassification of the less populated ones increased. In the case of ADHD-200 database, this led to useless classifiers typically selecting the most populated class.

**Table 7:**
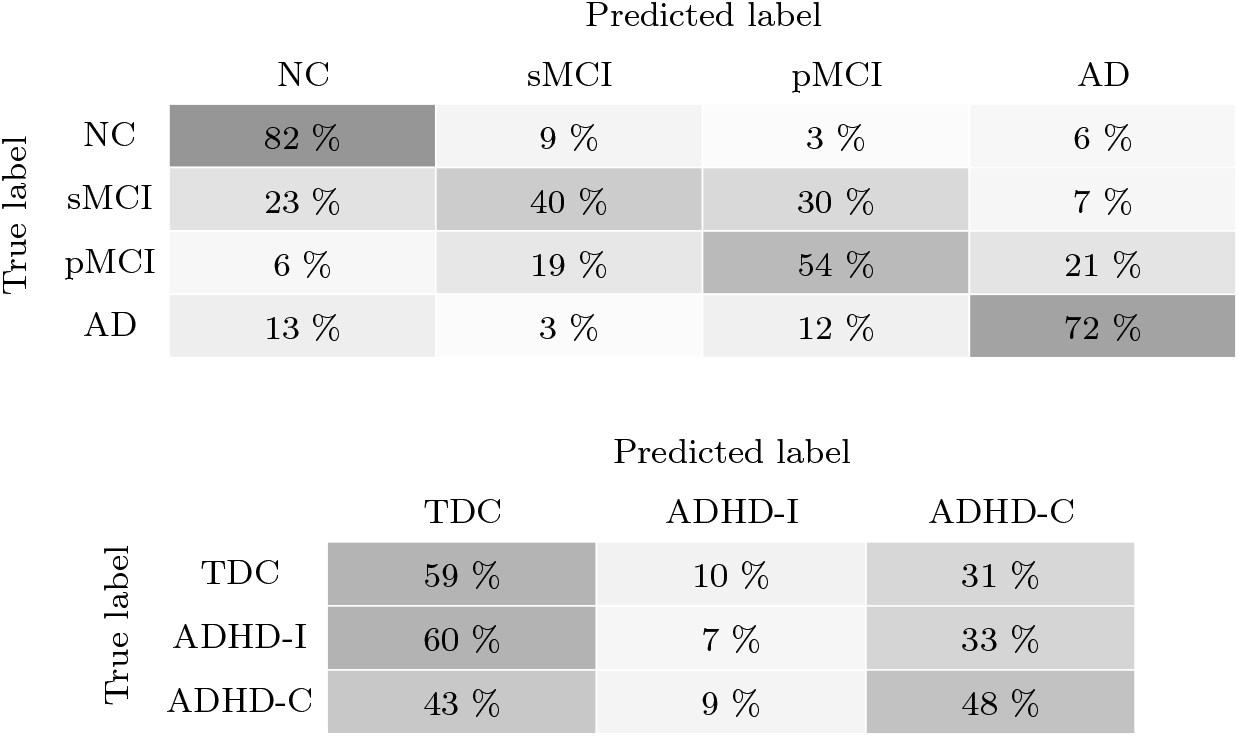
Classifier’s confusion matrix with the balanced RB-CCA+ST and validating with the balanced accuracy. With the ADNI database (left) the method is capable of correctly classify around 50% of the least populated classes while separating them from the most populated ones. With the ADHD database (right) the method provides a good classification of the highly less populated classes regarding the complexity of the problem.

|  |  | Predicted label |  |  |  |
| --- | --- | --- | --- | --- | --- |
|  |  | NC | sMCI | pMCI | AD |
| True label | NC | 82 % | 9 % | 3 % | 6 % |
|  | sMCI | 23 % | 40 % | 30 % | 7 % |
|  | pMCI | 6 % | 19 % | 54 % | 21 % |
|  | AD | 13 % | 3 % | 12 % | 72 % |

Table 7: Classifier’s confusion matrix with the balanced RB-CCA+ST and validating with the balanced accuracy.
|  |  | Predicted label |  |  |
| --- | --- | --- | --- | --- |
|  |  | TDC | ADHD-I | ADHD-C |
| True label | TDC | 59 % | 10 % | 31 % |
|  | ADHD-I | 60 % | 7 % | 33 % |
|  | ADHD-C | 43 % | 9 % | 48 % |

**Table 8:** Classifier’s confusion matrix with the balanced RB-CCA+ST and validating with the standard accuracy. With the ADNI database (left) the validation provides a greater misclassification of the least populated class (sMCI) while slightly improving the classification of the most populated ones. With the ADHD database (right) validating with the standard accuracy implies a substantial reduction of the classification of both least populated classes.

|  |  | Predicted label |  |  |  |
| --- | --- | --- | --- | --- | --- |
|  |  | NC | sMCI | pMCI | AD |
| True label | NC | 81 % | 7 % | 3 % | 9 % |
|  | sMCI | 27 % | 34 % | 33 % | 6 % |
|  | pMCI | 6 % | 21 % | 52 % | 21 % |
|  | AD | 13 % | 2 % | 11 % | 74 % |

Table 8: Classifier’s confusion matrix with the balanced RB-CCA+ST and validating with the standard accuracy.
|  |  | Predicted label |  |  |
| --- | --- | --- | --- | --- |
|  |  | TDC | ADHD-I | ADHD-C |
| True label | TDC | 87 % | 2 % | 11 % |
|  | ADHD-I | 86 % | 4 % | 10 % |
|  | ADHD-C | 79 % | 2 % | 19 % |

### Selected features and summary components

Figure 3 represents which voxels have been selected using RB-CCA+ST with ADNI data. Figure 3 shows the axial slices at locations that have been previously found to be to relevant AD-related cerebral atrophy including temporal, frontal and parietal areas, hypothalamus, cingulate gyrus and hippocampus (Weiner et al. 2017; Frisoni et al. 2010). As Figure 3 shows, the selections of the proposed method agreed with expectations based on literature. A more detailed figure of the class-wise selection including more axial slices is available in supplementary material.

**Fig. 3:**
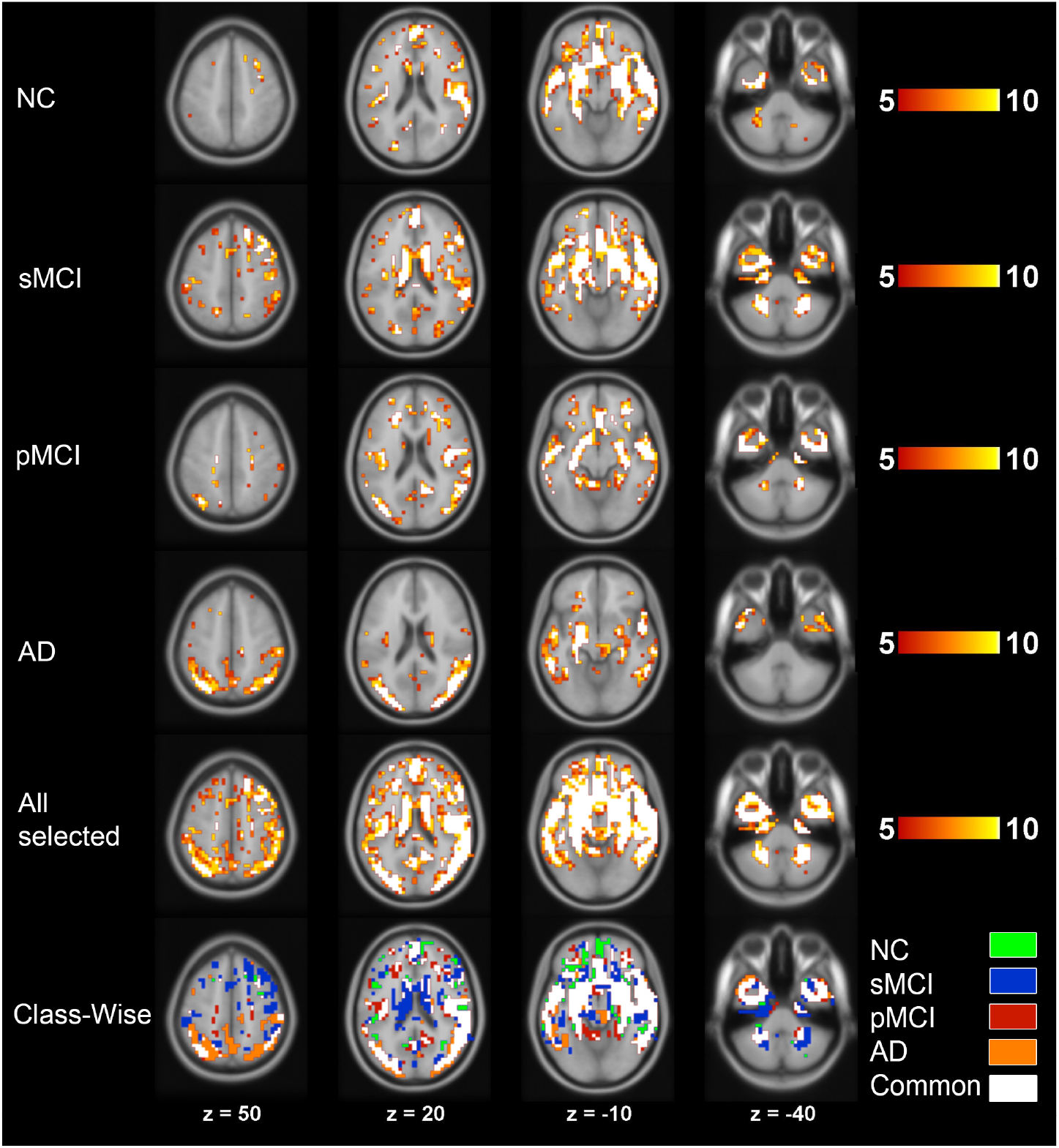
Locations of the most frequently selected voxels using RB-CCA with ADNI data. Note that the selection is first class-wise and each class-wise selected voxel is included in the final set of voxels. The overlay intensity gives the number of times a voxel has been selected during the 10-fold CV and we have used a threshold 5 to show only those voxels which have been selected in 50% of the folds. Four axial slices are shown, at *z* = 50*mm*, 20*mm* − 10*mm*, −40*mm* of the MNI space. The first 4 rows show the features selected for each class. The fifth row shows the complete selection which will be applied to the input data. The bottom row visualizes the classes providing the selected voxels.

Figure 4 shows the mean values of sign consistency parameters bjc over 10 CV-folds, note *b*_*jc*_ takes values between 0 and 1. Similarly to Figure 3, Figure 4 shows first class-wise sign consistencies, and the bottom row shows the combined relevance of all classes.

**Fig. 4:**
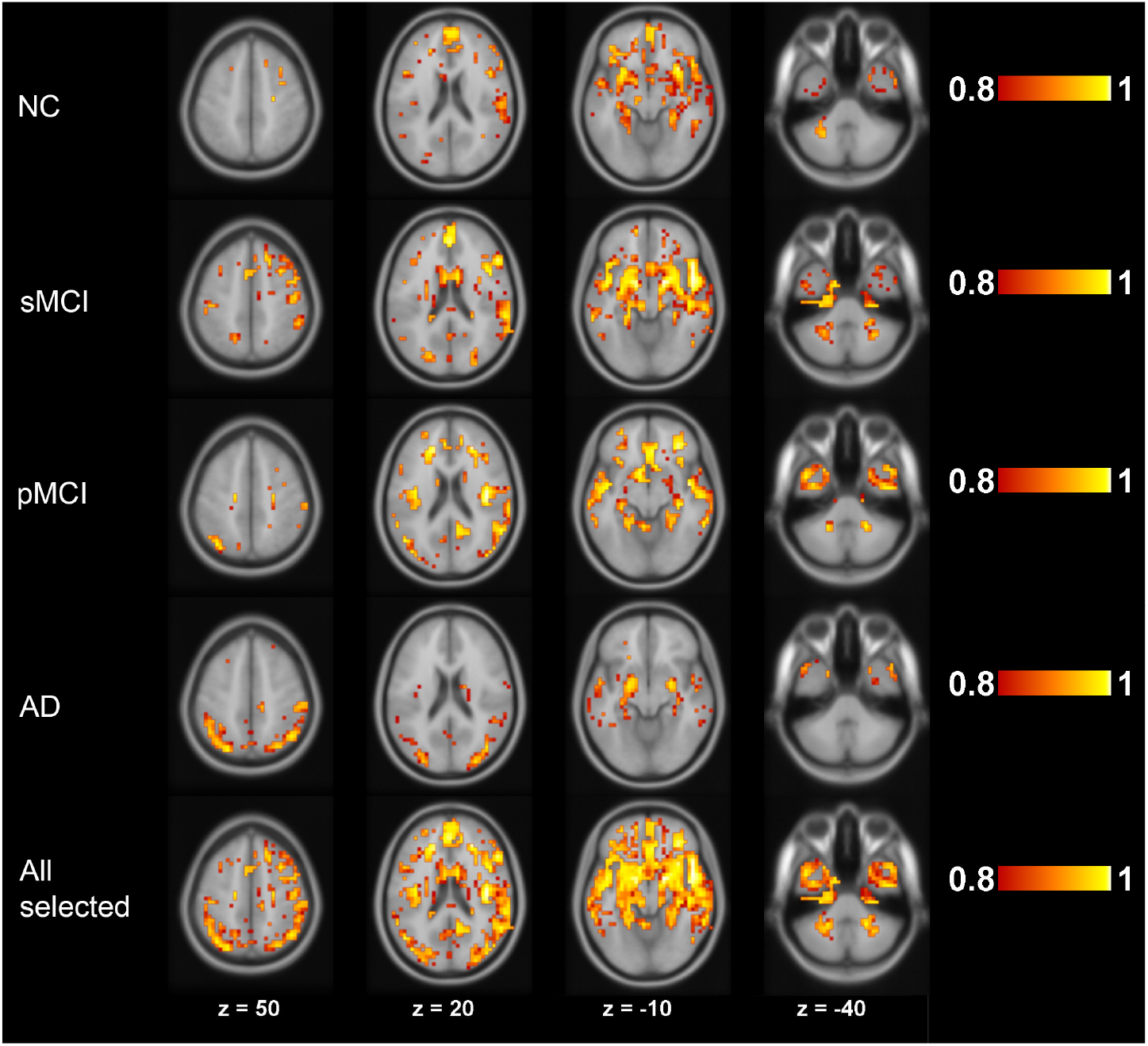
Variable relevance in ADNI data using RB-CCA. Only relevances of voxels which have been selected at least in 5 of 10 CV-folds are shown. Four axial slices are shown, at z = 50*mm*, 20*mm* − 10*mm*, −40*mm* of the MNI space. Four top rows show the class-wise relevances of the voxels and the bottom row shows the complete the relevance of all the selected voxels.

The summary components were generated by the CCA as a projection of the data in an orthogonal space that decorrelates the data (Equation (15)). In our four-class problem with ADNI data, three summary components were obtained using the information of the sign and magnitude voxel-wise consistency and were subsequently used by the classifier. In the feature extractor CCA (Equation (1)), the summary components are given by the primal projection matrix *U*, which shows the relation between the original features and the summary components. Figure 5 shows the first, second and third eigenvectors averaged over the 10 folds, which indicates how the selected voxels influence the construction of the three summary components. Figure 6 visualizes the summary components values for one representative CV fold. Figure 6a depicts the first two summary components along with the SVM boundaries between each pair of classes for a particular fold. The two summary components are decorrelated and orthogonal. The first summary component separated AD subjects from the rest and the second summary component separates the three other classes. Note that the class overlapping in Figure 6 is intuitive as pMCI subjects should be more similar to AD subjects and sMCI subjects should share a certain degree of similarity with NC subjects. Figure 6b shows the projection of the data using the three extracted summary components, which have been normalised for a better interpretation. The complete projection of the summary components is not as intuitive as its two dimensional form and the third component provides less help in the discrimination between the different classes.

**Fig. 5:**
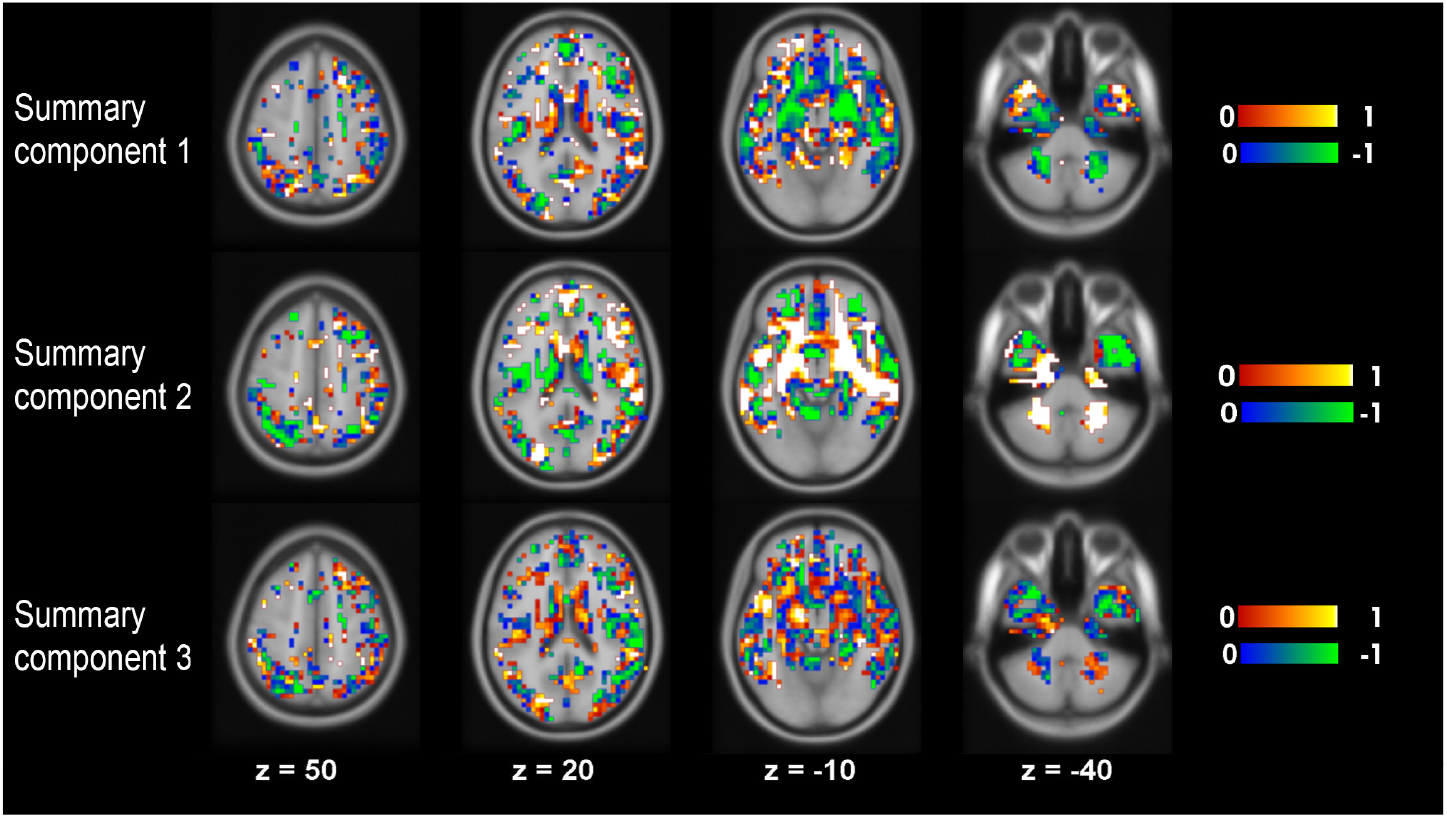
Normalised mean values of the generated summary components using RB-CCA with ADNI data. Masked with the most selected voxels in the 10-folds CV.

**Fig. 6:**
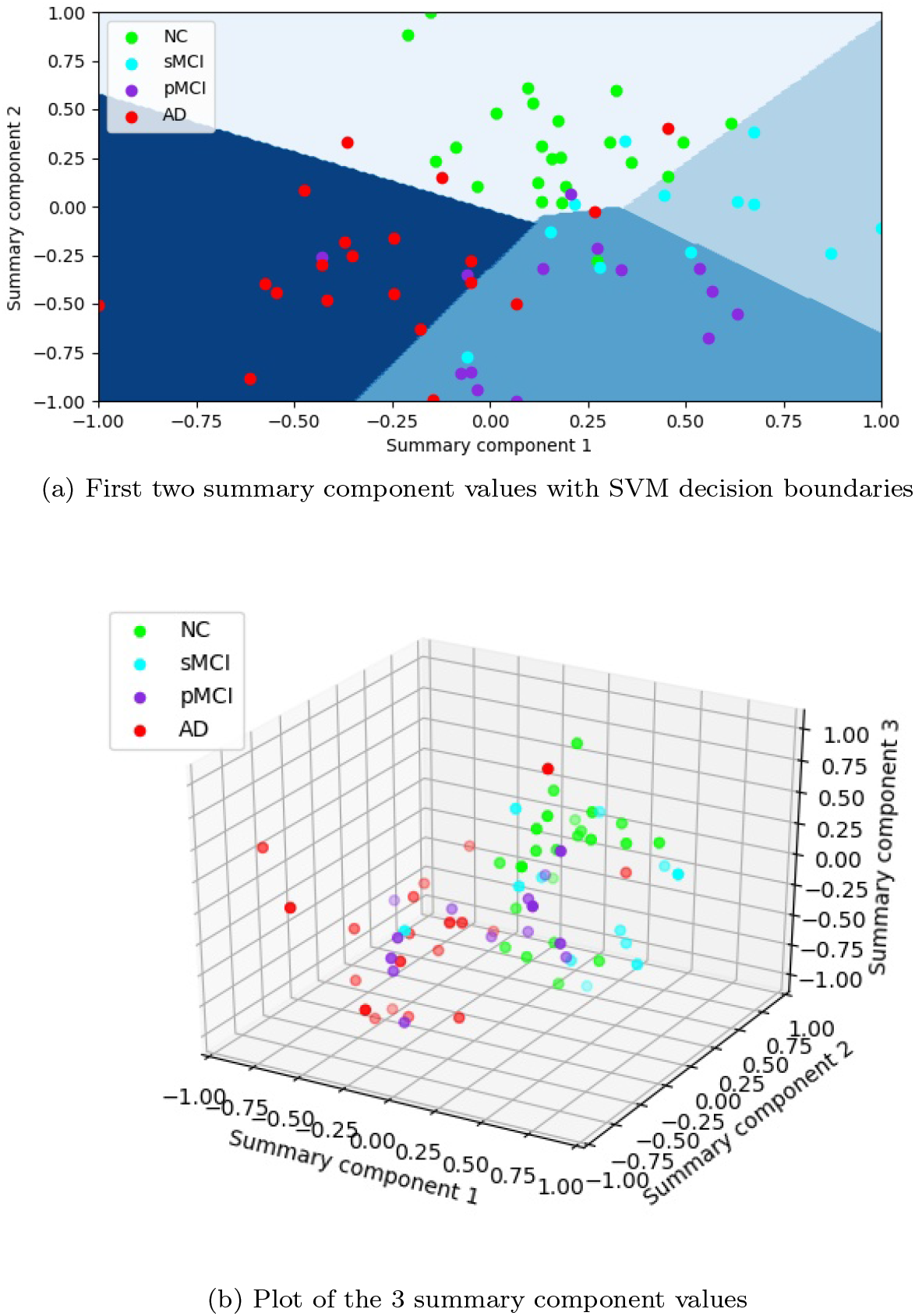
Normalised summary component values for one representative fold with the SVM’s decision boundaries using ADNI data. The proposed method finds a projection of the data capable of representing the dataset with 3 values. The first two summary components are the most informative and the ones that have the biggest role in the projection of the data. The first summary component is capable of discriminating between sMCI and AD. The second summary component separates NC from pMCI.

Regarding the ADHD database the same procedure has been followed as with the ADNI database. However, as this database is composed by fMRI, the plot of the data is less intuitive, complicating the interpretation of the results. For this reason, the images related to the selected features and the summary components have been included in the supplementary material.

For the ADHD dataset, the summary component values are shown in Figure 7, having the plot of the summary components along with the SVM boundaries between each pair of classes for a particular fold. As happened with the previous database, the main problem with this data resides on the discrimination of the minority classes. This can be seen in this figure, were the less populated classes are partially separated but do not provide any kind of conclusive result.

**Fig. 7:**
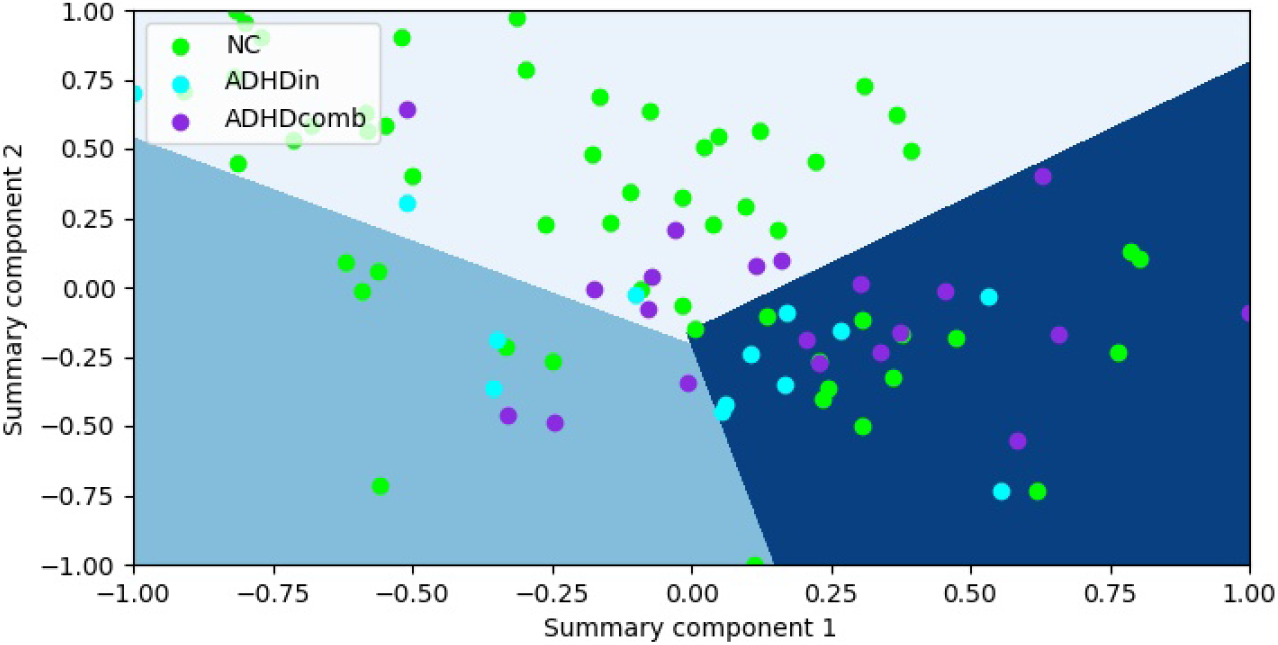
ADHD - Normalised summary components for one representative fold with the SVM’s boundaries. The proposed method finds a projection of the data capable of representing the dataset with 2 values.

## Discussion

This paper has presented a RB-CCA framework for the extraction of summary components in neuroimaging data. The method first carries out a bagging procedure, which calculates the sign consistency of the CCA projection matrix feature-wise, to determine the relevance of each feature. Then, it uses the learned relevance to select the most significant features and to regularize the posterior CCA. To select the optimum number of features, we have proposed a novel hypothesis test based approach as a replacement for the CV-based model selection, which is time consuming and prone to high variance of the CV based error estimates. The proposed method combines a FS and a FE step capable of using the information obtained in the FS process to guide, according to the importance of each feature, the subsequent FE stage. The final result of the method are the summary components which reduce the original number of features to just *c* − 1 orthogonal components that are easy to visualize and provide insights to the data.

The proposed method is inspired by Muñoz-Romero et al. (2017), but contains several novel aspects as compared to it: 1) We have adapted the bagging procedure to obtain a class-wise FS which improves the interpretability of the feature selection, having the features that are most relevant for each class. 2) The additional computational cost derived from this change is solved by the inclusion of a statistical hypothesis test, specifically designed for the bagged CCA scheme, to be able to automatically select the optimum number of selected features per class. The hypothesis test removes the necessity to carry our computationally expensive CV to select the optimum threshold for feature selection and we have shown that it leads to equal classification accuracy than the use of CV. 3) We have improved the regularized CCA by making it use the sign consistency, which is obtained through the bagged CCA, in its regularization term. This leads to novel regularization based in both sign and magnitude consistency. This modification provides a more informative regularization by the inclusion of extra relevance criterion that has been learned during the CCA bagging. 4) We have introduced the dual space formulation of CCA leading computational savings in small-sample high dimensional scenarios. The previous approach assumed that having regularized feature extractor was enough to use the primal formulation, which is advantageous when (*N* > *d*), where *d* refers to the number of selected features. However, when sample sizes are small enough compared to the data dimensionality, as often in brain imaging, the dual formulation is advantageous. 5) Finally, we have introduced balanced formulation of MVA to enable its use in heavily imbalanced problems as these are typical in brain imaging.

We have applied the different variations of the proposed method to two different databases. With the ADNI database using the proposed parsimonious MVA along with the feature selection and the ST provide the best results in terms of balanced accuracy and AUC. At the same time, the balanced version of the method outperforms the unbalanced when there is a dataset with highly unbalanced classes. Using this optimum set-up the method provides a balanced accuracy of 61, 56% in the multiclass classification setting, which poses an improvement of 5% in the balanced accuracy compared to the different analysed baselines and 0, 036 in terms of AUC.

The accuracy results obtained with our method in the ADNI database compared to the one presented by Liu et al. (2015) for the 4 classes classification, pose an improvement of almost 0,14 in the standard (not balanced) accuracy. Furthermore, we have implemented the method p-map used by Abdulkadir et al. (2014) in the CADDementia challenge (Bron et al. 2015). The comparison between our method and p-map showed an improvement of 0,07 in terms of accuracy and 0,03 in terms of AUC.

Our results suggested that in the ADNI database the sMCI class was the most difficult to classify. This matches with Dong et al. (2016) where different clusters where defined to differentiate MCI subjects. In this sense, the proposed balanced version was able to pay more attention over the sMCI class, facilitating the detection of this group of subjects.

The advantages were not as clear with the ADHD database as with the ADNI database. If we compare our performance to that of the baseline methods without feature selection (SVM and CCA), the results show that our method was capable of providing similar results in terms of balanced accuracy and slightly better in terms of AUC while using roughly 30% of the original variables.

In relation to the methods which carry out feature selection, we have compared to p-map (Liu et al. 2015) and RFE+HELM (Qureshi et al. 2016), which have been implemented and validated using the balanced accuracy measure. Note that we can not directly compare to the results obtained in Qureshi et al. (2016) as they were working with ROIs instead of voxels and they did not use the balanced accuracy to validate their method and to evaluate its performance. Our experiments showed that our method achieves an improvement of 0, 02 and 0, 09 in balanced accuracy and 0, 03 and 0, 07 in AUC, compared to p-map and RFE+HELM, respectively.

The balanced version of our method outperforms the unbalanced version, which overfits to the majority class. Most multiclass methods proposed in neuroimaging (e.g., (Qureshi et al. 2016, 2017)) deal with class imbalance by reducing the number of subjects of all classes to the number of subjects of the minority class. Our method tackles the class imbalance by the definition of a balanced version able to maintain all the available data.

While most methods for multiclass taks in neuroimaging are problem-specific (Bron et al. 2015; Qureshi et al. 2016), our approach is more generic. The method is able to work with very high dimensional data, and as shown in the experiments, overcomes the limitations by other methods designed to use fewer variables. Regardless of the limits imposed by the difficulty of the datasets, our method showed a good performance at the selection of most relevant features, the definition of summary components and classification. The summary components have proven to by sufficiently informative to describe with two or three of them the whole set of original variables. Finally, the scoring results show that the method is capable of working with the multiclass setting, which has not been widely studied, providing consistent results for the different scenarios analysed.

## Acknowledgments

Data collection and sharing for this project was funded by the Alzheimer’s Disease Neuroimaging Initiative (ADNI) (National Institutes of Health Grant U01 AG024904) and DOD ADNI (Department of Defense award number W81XWH-12-2-0012). ADNI is funded by the National Institute on Aging, the National Institute of Biomedical Imaging and Bioengineering, and through generous contributions from the following: AbbVie, Alzheimer’s Association; Alzheimer’s Drug Discovery Foundation; Araclon Biotech; BioClinica, Inc.; Biogen; Bristol-Myers Squibb Company; CereSpir, Inc.; Cogstate; Eisai Inc.; Elan Pharmaceuticals, Inc.; Eli Lilly and Company; EuroImmun; F. Hoffmann-La Roche Ltd and its affiliated company Genentech, Inc.; Fujirebio; GE Healthcare; IXICO Ltd.; Janssen Alzheimer Immunotherapy Research & Development, LLC.; Johnson & Johnson Pharmaceutical Research & Development LLC.; Lumosity; Lundbeck; Merck & Co., Inc.; Meso Scale Diagnostics, LLC.; NeuroRx Research; Neurotrack Technologies; Novartis Pharmaceuticals Corporation; Pfizer Inc.; Piramal Imaging; Servier; Takeda Pharmaceutical Company; and Transition Therapeutics. The Canadian Institutes of Health Research is providing funds to support ADNI clinical sites in Canada. Private sector contributions are facilitated by the Foundation for the National Institutes of Health (www.fnih.org). The grantee organization is the Northern California Institute for Research and Education, and the study is coordinated by the Alzheimer’s Therapeutic Research Institute at the University of Southern California. ADNI data are disseminated by the Laboratory for Neuro Imaging at the University of Southern California.

C. Sevilla-Salcedo and V. Gómez-Verdejo’s work has been partly funded by the Spanish MINECO grant TEC2014-52289R and TEC2017-83838-R. Jussi Tohka’s work is supported by the Academy of Finland (grant 316258).

Alzheimer’s Disease Neuroimaging Initiative (ADNI) is a Group/Institutional Author.

Data used in preparation of this article were obtained from the Alzheimer’s Disease Neuroimaging Initiative (ADNI) database (adni.loni.usc.edu). As such, the investigators within the ADNI contributed to the design and implementation of ADNI and/or provided data but did not participate in analysis or writing of this report. A complete listing of ADNI investigators can be found at: http://adni.loni.usc.edu/wp-content/uploads/how_to_apply/ADNI_Acknowledgement_List.pdf

## A Hypothesis Test

Considering the success probability *p_jcr_* = (1/*P*) 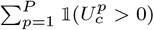, we can formulate the following hypothesis test:

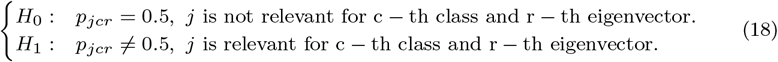

To be able to statically evaluate if *p*_*jcr*_ differs from 0.5, we define the following statistic:

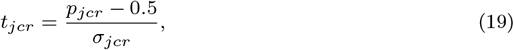

where *σ*_*jcr*_ is a scaling factor proportional to the standard error of *p*_*jcr*_. We now derive this scaling factor. The term 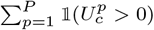 counts of the number of times that a feature is positive over *P* bagging iterations. Thus, assuming that the bagging iterations are independent, it can be modelled as a rescaled Binomial distribution with parameters *P* (number of experiments) and *p*_*jcr*_ (success probability). Further, since the number of bagging iterations is very large, the binomial distribution can be approximated by a Normal distribution with mean *P* · *p*_*jcr*_ and variance *P* · *p*_*jcr*_(1 − *p*_*jcr*_). So, under the independence assumption, we can define *σ*_*jcr*_ as the standard deviation of the term 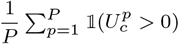, which is straightforwardly computed by rescaling the variance of the Normal distribution:

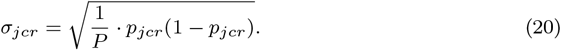

However, we need to take into account that the observations are coming from a bagging process and independence can not be assumed. To address this problem, the standard deviation is computed with an unbiased estimator (Nadeau and Bengio 2000) which, applied to our scenario, provides the following corrected estimator for the standard deviation:

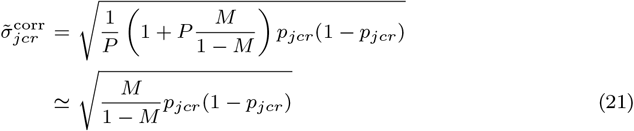

and, therefore, the statistic *t*_*j*_ becomes:

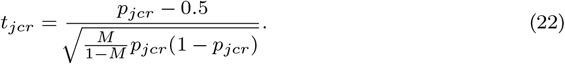

The statistic *t*_*jcr*_ is distributed according to the t-distribution with *P* − 1 degrees of freedom. Since *P* is very large, one can safely approximate the t-distribution by the standard normal distribution.

Once this statistic is calculated, the class-wise feature selection can be carried out by majority-voting r. This means that for each class we select the features that are considered as relevant by the majority of the eigenvectors.

Thanks to the inclusion of the statistical test, the cross-validation (CV) of the optimum amount of selected features is not needed, therefore reducing the computational time. Furthermore, this efficient approach allows the selection of features in a class-wise manner, improving the interpretability of the results and posing an advantage over the approach presented in Muñoz-Romero et al. (2017).

## B Unbalanced method results

In this appendix further results obtained with different versions of the method are depicted. In particular, here we present the results obtained in both databases when not using the balanced version of the method.

Regarding Table 9, the results are similar to the ones obtained in Table 5 in terms of accuracy and slightly worse considering the AUC. The main advantage of using the balanced version in this database is the improvement in the classification of the most critical class, sMCI.

**Table 9:**
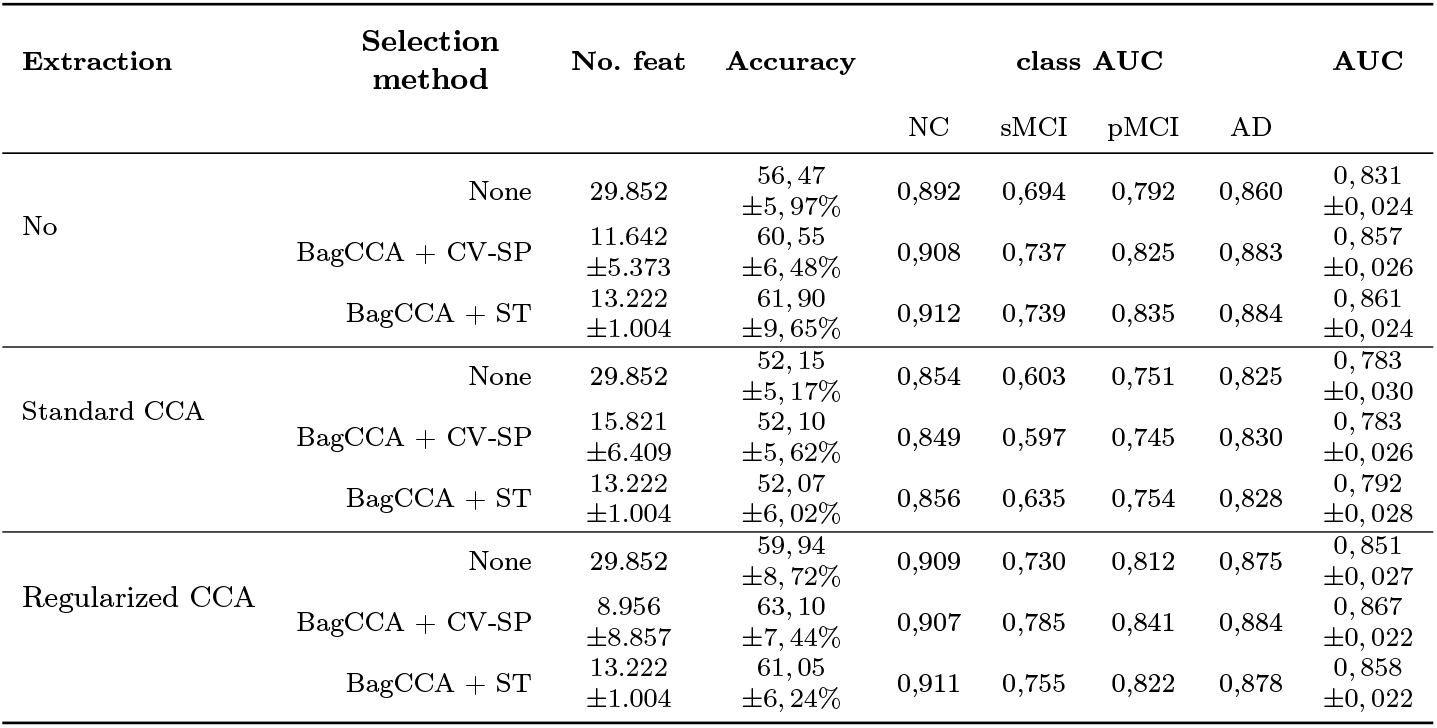
ADNI - Accuracy results with the different versions of the method, considering the usage of the proposed selection and extraction methods in their unbalanced version. This table justifies the need of adding the regularisation to the CCA as well as the usage of a selection method. Furthermore, this table depicts the need to include the balanced version for the correct classification of less populated classes.

In Table 10 we can see the results obtained using the unbalanced version of the method in the ADHD database. In this highly unbalanced database, the results are worse than the ones obtained in Table 6 in terms of accuracy, having that without the usage of the balanced version the method overfits to the most populated class. Therefore, it is critical in this database to use the balanced version.

**Table 10:**
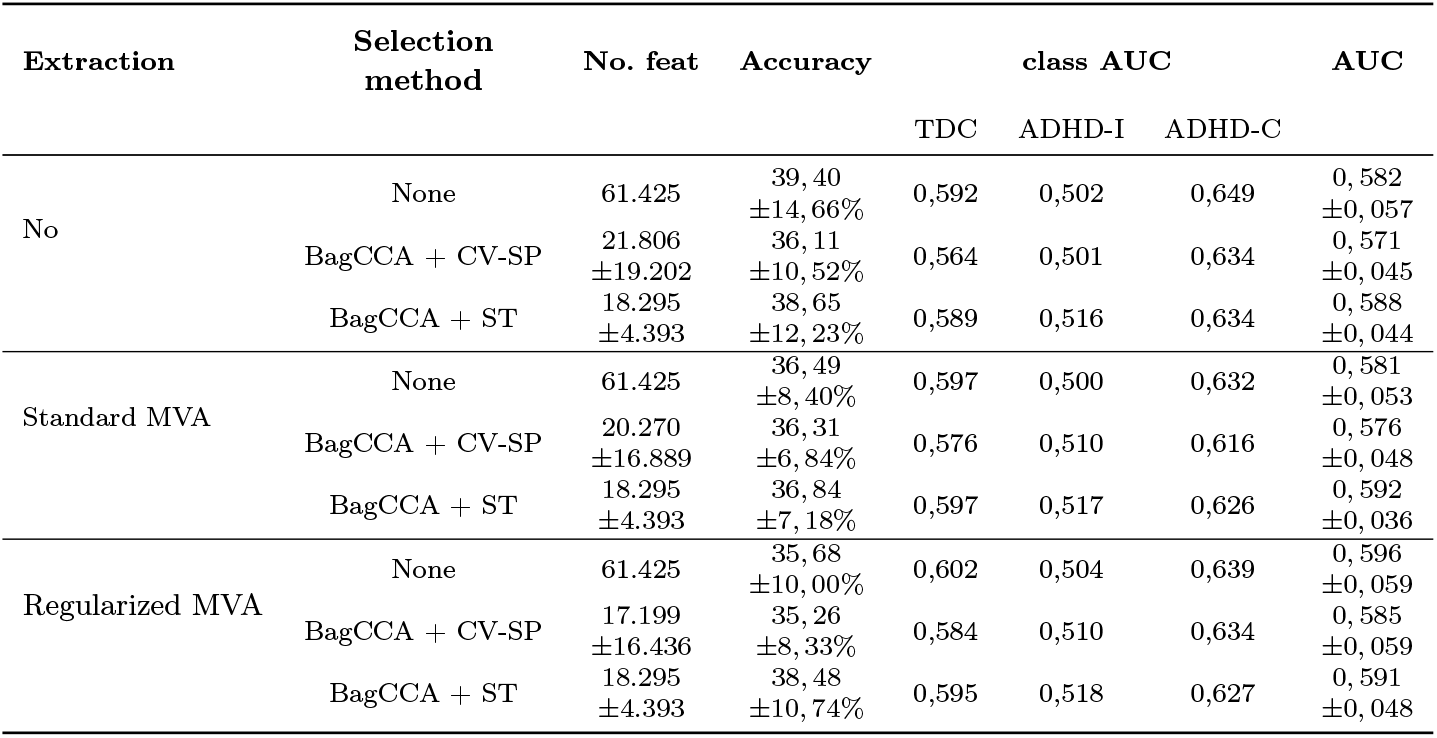
ADHD - Accuracy results with the different versions of the method, considering the usage of the proposed selection and extraction methods in their unbalanced version. The feature selection improves the interpretability of the results, reducing them by one fifth while keeping a similar performance. Nevertheless, the results obtained without the balanced version imply overfitting the most populated class and do not provide reliable results.

## Footnotes

1 The reference Bron et al. (2015) summarizes the results of the data analysis competition, where the task was to classify subjects into Alzheimer’s disease (AD), mild cognitive impairment (MCI), and cognitively normal classes. We use this summary paper as a reference to all the methods in the competition if there is no particular reason to specify a particular method.

2 Note that the inclusion of the regularization term over A prevents problems in the calculation of the inverse of *K*_*x*_*K*_*x*_. These issues should not appear when working with high dimensional data, however they can occur in case of high redundancy among variables.

3 As the accuracy validation curves tend to present a saturation profile and their maximum value is given when almost all features are used, we have selected as optimum working point the CV Stability Point (CV-SP), the point of the curve where the saturation begins. In this way, we obtain a good performance point using a reduced set of features.

