## Supplementary Material for "Regularized Bagged Canonical Component Analysis for Multiclass Learning in Brain Imaging"

**Neuroinformatics manuscript No.**
(will be inserted by the editor)

### Regularized Bagged Canonical Component Analysis for Multiclass Learning in Brain Imaging - Supplementary material

**Carlos Sevilla-Salcedo · Vanessa Gómez-Verdejo · Jussi Tohka · Alzheimer's Disease Neuroimaging Initiative**

Received: date / Accepted: date

In this supplementary material we include some of the layouts that were too extensive to include in the paper and can provide a better insight of the analysis developed.

First of all, in relation to the ADNI database, we have included an image with the class-wise feature selection with more slices than the 4 that were included in the article. Therefore, in Figure 1 it can be seen that there are different brain areas that are predominant for a specific class. This way the fourth row distinctively specifies some areas that are selected for AD and sMCI, while in the seventh and eighth row there are some relatively wide regions of voxels significant for NC and pMCI.

In relation to the ADHD database, we have included the results obtained using the proposed method. These images show the complete correlation matrices obtained from the 351 regional time courses, as explained in the paper. In particular, with these images we show the effect of the method on the input features.

Figures 2, 3 and 4 show the different measures obtained for the features of the database. Unlike ADNI, the features of this database are not easily interpretable. The main conclusion obtained from these images is that there is a feature selection that do not worsen the results.

Carlos Sevilla-Salcedo
Department of Signal Processing and Communications, Universidad Carlos III de Madrid, Leganés, Spain

ORCID: 0000-0002-5507-7537

Vanessa Gómez-Verdejo
Department of Signal Processing and Communications, Universidad Carlos III de Madrid, Leganés, Spain

Jussi Tohka
A.I. Virtanen Institute for Molecular Sciences, University of Eastern Finland, Kuopio, Finland

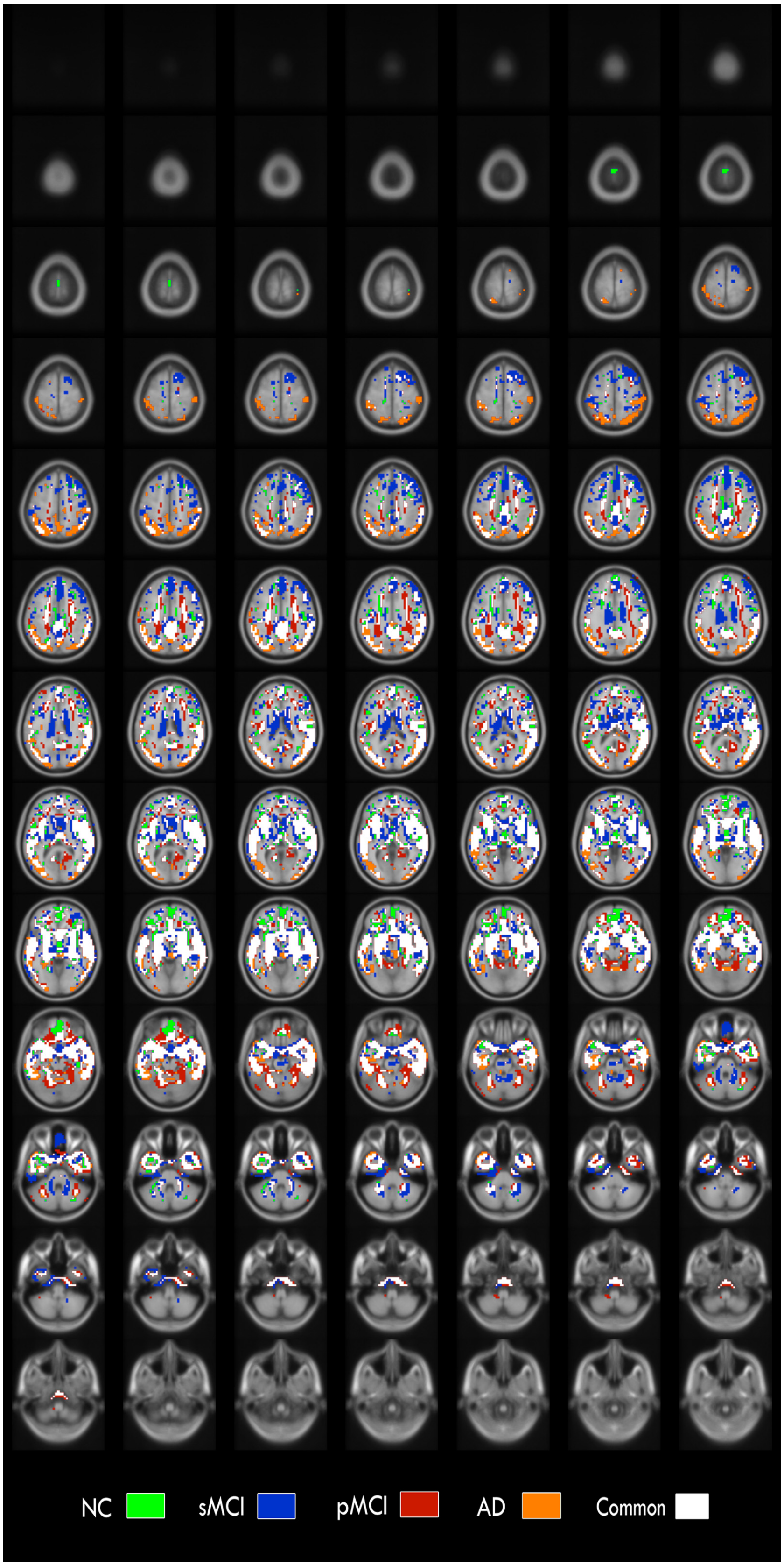


Fig. 1: Locations of the most frequently selected voxels using BagCCA + ST with ADNI data. The selection is class-wise selected voxel included in the final set of voxels. The overlay intensity gives the number of times a voxel has been selected during the 10-fold CV and we have used a threshold 5 to show only those voxels which have been selected in 50% of the folds. 91 axial slices are shown, from $z = 108mm$ to $z = -72mm$ with steps of $2mm$ of the MNI space.

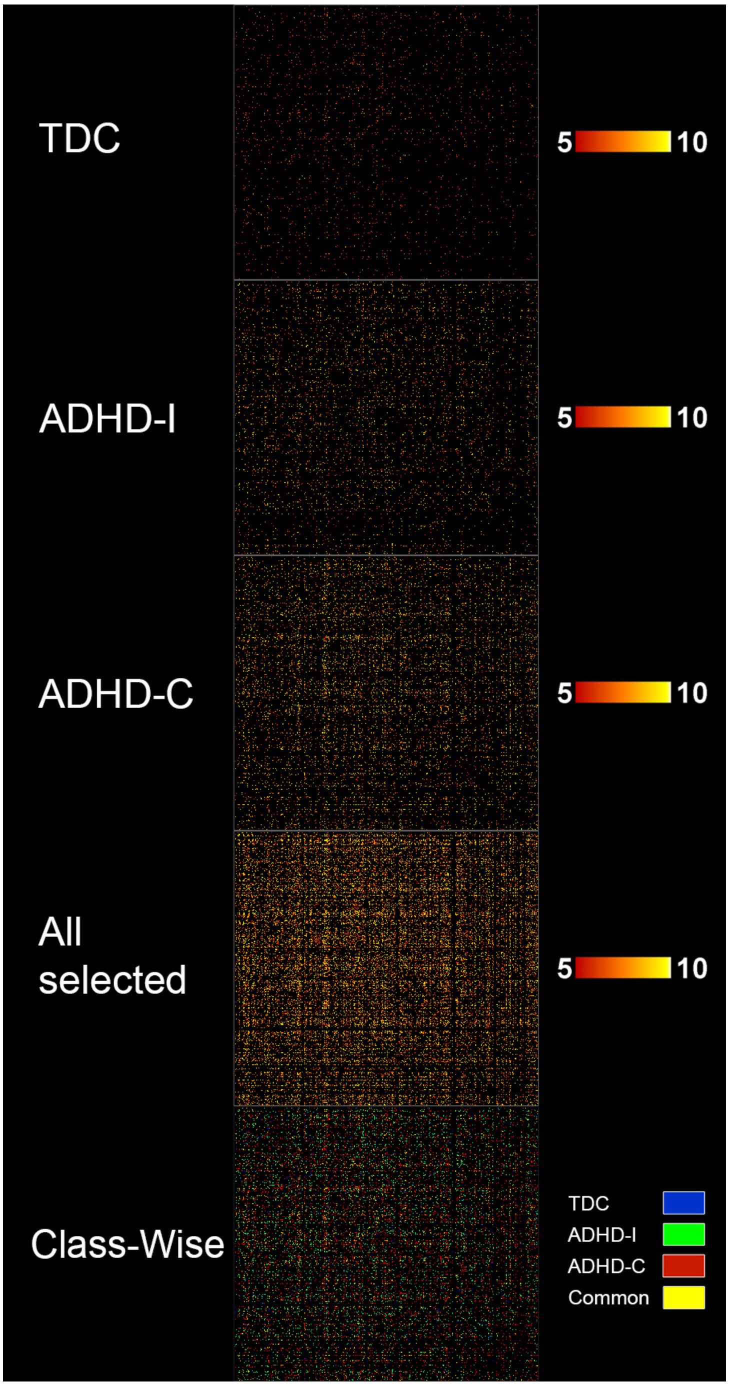


Fig. 2: Locations of the most frequently selected features using BagCCA + ST with ADHD data. The image depicts the complete correlation matrices obtained from the 351 regional time courses. The selection is class-wise selected features included in the final set of features. The overlay intensity gives the number of times a feature has been selected during the 10-fold CV and we have used a threshold 5 to show only those features which have been selected in 50% of the folds. The first 3 rows show the features selected for each class. The fourth row shows the complete selection which will be applied to the input data. The bottom row visualizes the classes providing the selected features.

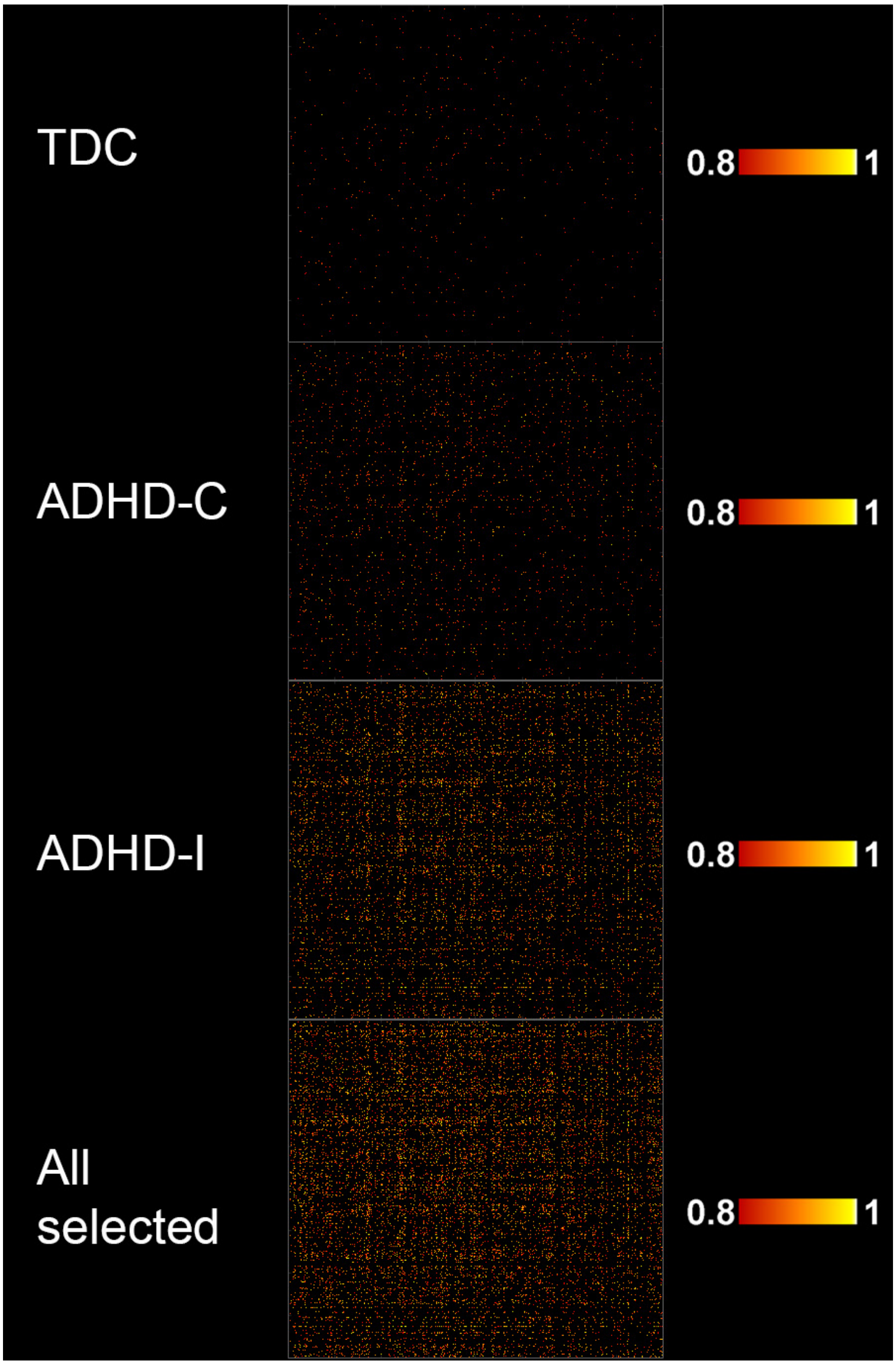


Fig. 3: Variable relevance using BagCCA + ST with ADHD data. The image depicts the complete correlation matrices obtained from the 351 regional time courses. Only relevances of features which have been selected at least in 5 of 10 CV-folds are shown. Three top rows show the class-wise relevances of the features and the bottom row shows the complete the relevance of all the selected features.

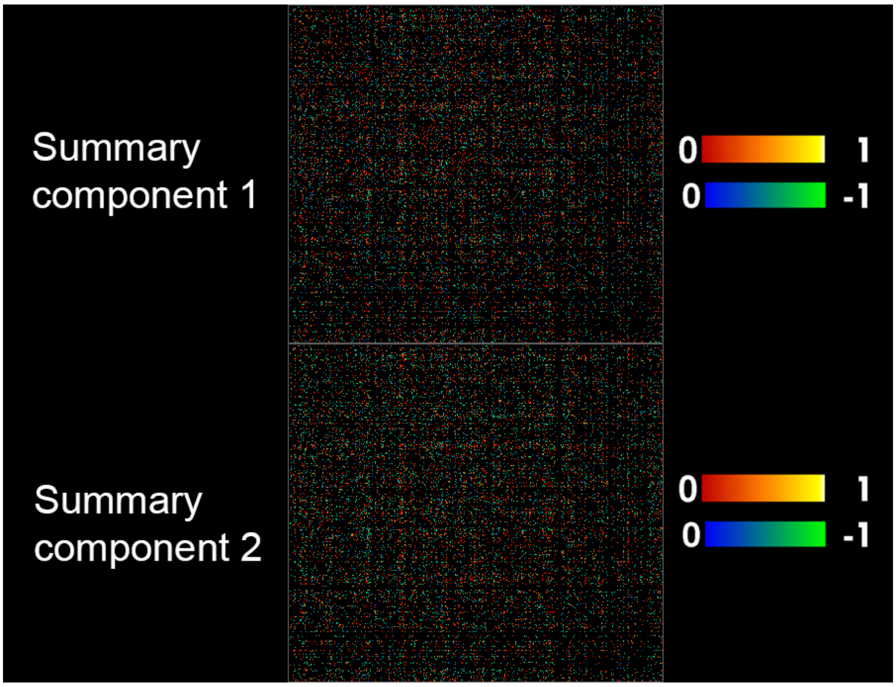


Fig. 4: Normalised mean values of the generated summary components using BagCCA + ST with ADHD data. The image depicts the complete correlation matrices obtained from the 351 regional time courses. Masked with the most selected features in the 10-folds CV.
